# *Ca*Pti1/*Ca*ERF4 module positively regulates pepper resistance to bacterial wilt disease through coupling enhanced immunity and dehydration tolerance

**DOI:** 10.1101/2021.10.26.465959

**Authors:** Lanping Shi, Xia Li, Yahong Weng, Hanyang Cai, Kaisheng Liu, Baixue Xie, Hussain Ansar, Deyi Guan, Shuilin He, Zhiqin Liu

## Abstract

Bacterial wilt, a severe disease that affects over 250 plant species, is caused by *Ralstonia solanacearum* through vascular system blockade. Although both plant immunity and dehydration tolerance might contribute to disease resistance, whether and how they are related are still unclear. Herein, we provide evidence that immunity against *R. solanacearum* and dehydration tolerance are coupled and regulated by *Ca*Pti1-*Ca*ERF4 module. By expression profiling, virus-induced gene silencing in pepper and overexpression in *Nicotiana benthamiana*, both *CaPti1* and *CaERF4* were upregulated by *R. solanacearum* inoculation, dehydration stress and exogenously applied ABA. They in turn phenocopied with each other in promoting pepper resistance to bacterial wilt not only by activating HR cell death and SA-dependent *CaPR1*, but also by activating dehydration tolerance related *CaOSM1* and *CaOSR1*, and stomata closure to reduce water loss in ABA signaling dependent manner. Yeast-two hybrid assay showed that *Ca*ERF4 interacts with *Ca*Pti1, which was confirmed by co-immunoprecipitation and pull-down assays. Chromatin immunoprecipitation and electrophoretic mobility shift assay showed that, upon *R. solanacearum* inoculation, *CaPR1, CaOSM1* and *CaOSR1* were directly targeted and positively regulated by *Ca*ERF4 via binding GCC-box or DRE-box, which was potentiated by *Ca*Pti1. In addition, our data indicate that *Ca*Pti1-*Ca*ERF4 complex might act downstream ABA signaling, since the exogenously applied ABA did not alter stomata aperture regulated by *Ca*Pti1-*Ca*ERF4 module. Importantly, *Ca*Pti1-*Ca*ERF4 module was found also acts positively in pepper growth and response to dehydration stress. Collectively, the results suggest that immunity and dehydration tolerance are coupled and positively regulated by *Ca*Pti1-*Ca*ERF4 in pepper plants to enhance resistance against *R. solanacearum*.

**Summary:** Pepper immunity and dehydration tolerance are coupled and regulated by the *Ca*Pti1-*Ca*ERF4 module partially in a way related to ABA signaling and stomata closure.

## Introduction

In their natural habitats, plants are frequently encountered by pathogens with various lifestyles and invasion strategies. To protect themselves from the attacks of pathogens, plants have to activate appropriate defense responses after perceiving the pathogens by massive transcriptional reprogramming with the action of various transcription factors (TFs). However, how the upstream signaling translated into appropriate defense responses against different kinds of pathogens by these TFs is not fully understood.

It has been well established that plants employ two types of receptors, cell surface localized PRRs and intracellular R-proteins, to detect and bind pathogen associated molecular patterns (PAMPs) and effectors to trigger PAMP triggered immunity (PTI) and effector triggered immunity (ETI), respectively. ETI, which is generally coupled with hypersensitive response (HR), is more robust, intensive and longer time prolonged than PTI, and is crucial in incompatible plant-pathogen interaction. However, accumulating evidence show that PTI and ETI share a number of signaling components (Lu and Tsuda, 2021). The common shared components might include Ca2+ signaling, MAPK cascades, reactive oxygen species (ROS), defense hormones including salicylic acid (SA), jasmonic acid (JA) and ethylene (ET), and abscisic acid (ABA). Among these components, the function of ABA in plant immunity appear to be contradictory, for it has been found to act negatively in plant immunity by antagonizing SA signaling pathway (de Torres-Zabala et al., 2007; de Torres Zabala et al., 2009; Moeder et al., 2010; Robert-Seilaniantz et al., 2011). Pto (*Pseudomonas syringae* pv. *tomato*) was the first R protein isolated from *Pseudomonas syringae-* infected tomato (Ronald et al., 1992; Martin et al., 1993). It encodes a Serine/Threonine (Ser/Thr) protein kinase that recognizes effector AvrPto or AvrPtoB secreted by *Pseudomonas syringae* (Tang et al., 1996; Kim et al., 2002; Pedley and Martin, 2003), to trigger immunity against *Pseudomonas syringae*. The signaling imitated via perception of AvrPto or AvrPtoB by Pto can be transmitted downstream through phosphorylating other serine/threonine kinases such as Pti1 and PTI1-2 (Zhou et al., 1995) or transcription factors (*Sl*Pti4, *Sl*Pti5, *Sl*Pti6) in the nuclei (Gu et al., 2002). The plasma membrane localized *Os*Pti1 was found be essential for its regulatory role in immune responses (Matsui et al., 2015). Interestingly, Pti1 has also been found to be related to other biological processes acting as nodes for crosstalk between plant immunity and other processes. For instance, the *ZmPti1* (*Zea mays*), which was induced by SA treatment and pathogenic infection, positively regulates dehydration and salt tolerance in maize (Herrmann et al., 2006). However, whether it is also the case in other plant species and how it fulfills such kind of function are currently unclear.

AP2/ERF constitutes one of largest plant transcription factor families (Pieterse et al., 2009; Broekgaarden et al., 2015; Huang et al., 2016). A total of 147 and 123 AP2/ERF proteins have been found in *Arabidopsis* and pepper, respectively, which can be structurally divided into three subfamilies, namely, AP2, RAV, and ERF. Some members in ERF subfamily function in plant growth (Xie et al., 2019) and development (Pei et al., 2021), but the majority of members in this subfamily have been implicated in responses of plants to biotic and abiotic stresses (Pre et al., 2008; Meng et al., 2013; Dong et al., 2015; Froschel et al., 2019) by binding GCC- or DRE-box in promoters of their target genes (Fujimoto et al., 2000; Chakravarthy et al., 2003; Tournier et al., 2003; Liu et al., 2006; Van der Does et al., 2013). For instance, ERF1, ERF2, ERF14, ORA59, ERF5, ERF6, and ERF9 positively or negatively regulate immunity against pathogens in *Arabidopsis* in a collaborative manner (Gu et al., 2002; Yi et al., 2004; Maruyama et al., 2013; Meng et al., 2013; Cao et al., 2015; Li et al., 2015; Huang et al., 2016); some other AP2/ERF TFs have been found to be involved in plant response to abiotic stresses such as drought stress. For example, TINY in *Arabidopsis* (Xie et al., 2019) and *Sl*ERF84 in tomato (Li et al., 2018) have been found to positively and negatively regulate dehydration responses, respectively. In addition, it has been found that a single ERF might be involved in more than one biological process and acting as convergent nodes for multiple biological processes (Jisha et al., 2015; Papdi et al., 2015), reflecting the complex functions of AP2/ERF TFs. However, how these TFs are activated by upstream signaling to activate appropriate defense response to the attacks of pathogens is not very clear yet.

*Ralstonia solanacearum*, a soil-borne bacterial pathogen, causes bacterial wilt disease in over 250 plant species (Mansfield et al., 2012; Jiang et al., 2017), including the agriculturally important solanaceaes including pepper (*Capsicum annuum*) and tomato (*Solanum lycopersicum*). *R. solanacearum* exclusively infects plants through the roots; it colonizes the root cortex to reach the vasculature and spread through xylem vessels (Mansfield et al., 2012), and eventually causes plant wilting by blocking water uptake. Recently, Gerlin *et al* found that the plant water loss observed during *R. solanacearum* infection was in a similar range to what was observed during drought on tomato, supporting that bacterial wilt can impose significant drought stress on the plant (Montesinos-Pereira et al., 2014; Gerlin et al., 2021). Both plant immunity and dehydration tolerance might contribute to the disease resistance, but whether and how they are coupled are still unclear. In the present study, using pepper as an example, we provided the evidence that *Ca*Pti1 and *Ca*ERF4 were upregulated by *R. solanacearum* inoculation, they in turn promoted pepper resistance to bacterial wilt not only by activating HR cell death and SA-dependent *CaPR1*, but also by activating dehydration tolerance related genes including *CaOSM1* and *CaOSR1* and closing stomata to reduce water loss in a ABA signaling dependent manner. Our result might provide a new perspective for deciphering the mechanisms underlying plant resistance to bacterial wilt and its control.

## Results

### *R. solanacearum* inoculation, exogenous ABA application, and dehydration stress induce *CaPti1*

In our unpublished transcriptome deep-sequencing (RNA-seq) data in pepper plants challenged with *R. solanacearum* inoculation (RSI) by root irrigation, an upregulated differentially expressed gene (DEG) encoding a TyrKc domain-containing protein arouse our interest. Deduced amino acid sequence of the DEG is 400 amino acids in length and its predicted pI is 8.09. By phylogenetic analysis, it shares high sequence identities to TyrKc domain-containing proteins from other species, including *Na*Pti1 (85%) and *Nt*Pti1 in tobacco (94%), *At*Pti1 in Arabidopsis (85%), *St*Pti1 in potato (87%), *Sl*Pti1 in tomato (87%), and *Zj*Pti1 in jujube (86%); we named it as *Ca*Pti1 because it shares the highest sequence similarity to the *At*Pti among all of TyrKc domain-containing proteins in *Arabidopsis thaliana* (Supplemental Fig. S1).

*CaPti1* expression was assayed by quantitative RT-PCR to confirm the data of RNA-seq and to assess its possible involvement in response to *R. solanacearum* in pepper plant. The results showed that *CaPti1* was upregulated by *R. solanacearum* at 3 hours post inoculation (hereafter hpi) and lasted until 24 hpi (Supplemental Fig. S2A). In addition, to assess its possible involvement in pepper response to abiotic stress, the responses of *CaPti1* to exogenous application of ABA related to abiotic stress response or drought stress were assayed (Supplemental Fig. S2, B and C); the result showed that *CaPti1* was upregulated by exogenous applied ABA as well as by drought. As plant tolerance to drought and plant immunity are usually stomatal related, the expression of *CaPti1* in guard cell of pepper leaves was also assessed by isolating guard cells from *R. solanacearum*-inoculated pepper plants. The result showed that *CaPti1* was significantly induced in guard cells and mesophyll cells in response to *R. solanacearum* inoculation (Supplemental Fig. S2D). All these data imply that *Ca*Pti1 might play a role in pepper immunity against *R. solanacearum* and drought tolerance in a way related to stomata regulation.

### Subcellular localization of *Ca*Pti1 in *N. benthamiana* plants

By agroinfiltration based method, the subcellular localization of *Ca*Pti1 was investigated in leaves of *N. benthamiana* by fusing to GFP (green fluorescent protein) driven by the constitutive *CaMV35S* promoter or its native promoter (Supplemental Fig. S3, A-D). The success of protein biosynthesis was confirmed by immunoblotting analysis (Supplemental Fig. S3, B and D). The result showed that when driven by *CaMV35S* promoter, *Ca*Pti1-GFP located in multiple subcellular compartments, including cytoplasm and nuclei in epidermal cells of *N. benthamiana* (Supplemental Fig. S3A). However, when driven by its native promoter, *Ca*Pti1-GFP protein accumulated in the membrane and guard cell upon *R. solanacearum*, while no fluorescent signal was detected in the controls (Supplemental Fig. S3C). The results indicate that *Ca*Pti1 was upregulated by *R. solanacearum* and located in the membrane and guard cells.

### *CaPti1* silencing significantly increased the susceptibility of pepper plants to *R. solanacearum*

To assay the role of *Ca*Pti1 in pepper immunity against *R. solanacearum*, we employed knock down assay using a approach of virus-induced gene silencing (VIGS) (Liu et al., 2002). To avoid the possible of off-target, a specific fragment within 3’ UTR of around 300 bps in length was used to construct the vector for *CaPti1* silencing. The specificity and efficiency of this silencing was assessed by measuring the transcript of *CaPti1-like* (*CaPti1* homologue in pepper) by RT-qPCR (Supplemental Fig. S4A). The result showed that the transcript level of *CaPti1-like* was not affected, but that of *CaPti1* was significantly reduced in the *TRV:CaPti1* pepper plants with or without *R. solanacearum* challenge (Supplemental Fig. S4A). Noticeably, the *TRV:CaPti1* pepper plants exhibited a phenotypes, such as dwarf, with fewer leaves and fewer roots than the wild type plants (Supplemental Fig. S4B). By challenging with *R. solanacearum, TRV:CaPti1* pepper plants were found to exhibit an accelerated and more serious wilt symptoms than the control plants at 7 days post-inoculation (Fig. 1, A and B), and higher disease indices from 7 to 13 dpi. In parallel, an increase in *R. solanacearum* growth was also found in TRV:*CaPti1* pepper plants compared with the control plants (Fig. 1, C and D). In addition, the transcript levels of defense-associated marker genes were significantly decreased in *CaPti1*-silenced pepper leaves than those in control pepper leaves at 24 hpi (Fig. 1E). The marker genes included *CaPR1* (Choi and Hwang, 2011), *CaHIR1* (Jung et al., 2008), and *CaPO2* (Choi and Hwang, 2012). Interestingly, two dehydration stress-associated genes, including *CaOSM1* (Lee et al., 2010; Choi du et al., 2013) and *CaOSR1* (Park et al., 2016), were upregulated by *R. solanacearum* systematically in leaves but not in the *R. solanacearum*-infected roots (Supplemental Fig. S5A, S5B). However, *CaPti1*-silencing significantly reduced the upregulation of the tested marker genes by *R. solanacearum* in the leaves (Supplemental Fig. S5A, S5B).These results indicate that the silencing of *CaPti1* reduced pepper immunity against *R. solanacearum* partially by downregulating dehydration stress-associated genes such as *CaOSM1* and *CaOSR1*.

**Figure 1.**
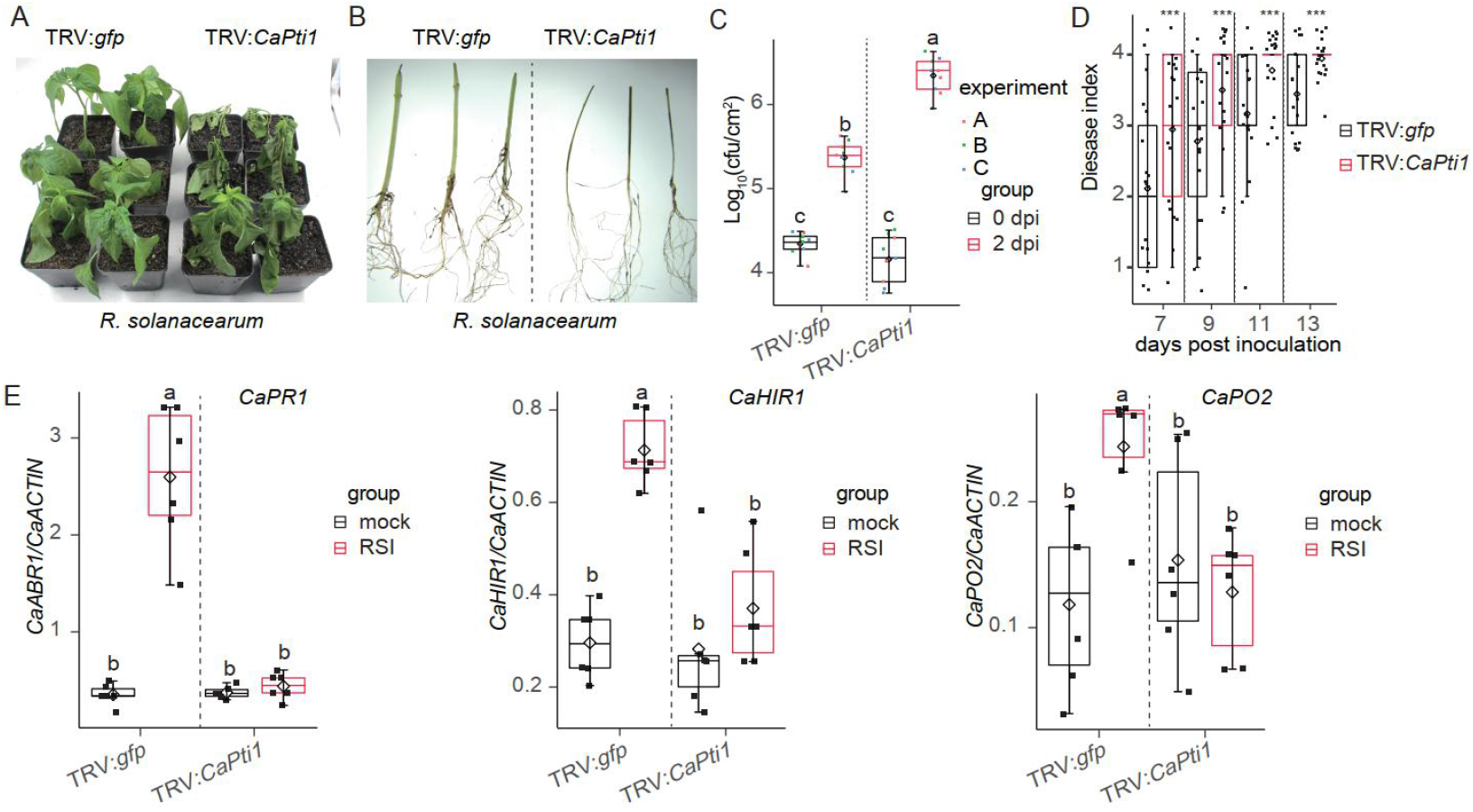
Silencing of *CaPti1* enhanced susceptibility of pepper plants to *R. solanacearum*. A, Phenotype of *CaPti1*-silenced and control pepper plants inoculated at the root with *R. solanacearum* (10^8^cfu/ml). Photographs were taken at 7 dpi. B, The phenotype of stems and roots in *CaPti1*-silenced and control pepper plants at 7 dpi. C, Growth of *R. solanacearum* in *R. solanacearum* inoculated *CaPti1*-silenced and control pepper plants at 2 hpi and at 2 dpi. Dots with different colors in boxplots represent 3 independent experiments with 6 biological replicates in each experiment (*n*=18). D, Progression of bacterial wilt in *CaPti1*-silenced and control pepper plants inoculated with *R. solanacearum* from 7 to 13 dpi. The data point (*n*=18) are present in boxplots. Asterisks indicate statistically significant differences as determined by the two-tailed *t* test (\*\*\**p*<0.001). E, The transcript levels of defense-relative marker genes, including *CaPR1, CaHIR1, CaPO2*, in *R. solanacearum* inoculated *CaPti1*-silenced and control pepper plants at 2 dpi. The mRNA expression levels of indicated genes were normalized to those of *CaACTIN*. Dots with different colors in boxplots represent 6 biological replicates (*n*=6). Three technical replicates for each biological sample were used. (C and E) Different letters above the bars indicate significant differences, as analyzed by Fisher’s protected LSD test (*p*<0.05).

### Ectopic overexpression of *CaPti1* enhanced resistance of *N. benthamiana* plants against *R. solanacearum*

We also evaluated the role of *Ca*Pti1 in immunity against *R. solanacearum* by gain-of-function assay. *CaPti1* overexpressing *N. benthamiana* plants were generated via *Agrobacterium-mediated* transgenic method. Over 10 independent transgenic *N. benthamiana* T3 lines with ectopic overexpression of *CaPti1* were obtained. The transcript levels of *CaPti1* in *CaPti1*-OE (overexpression) lines and wild type (WT) were confirmed by a quantitative RT-PCR. Two lines (line 2 and line 5) with middle levels of *CaPti1* transcript but without any growth defects were selected for further analysis (Fig. 2A). FJC100301, a highly virulent strain of *R. solanacearum*, was used to inoculate the *CaPti1-OE* lines and WT plants. The plants of *CaPti*-OE *N. benthamiana* lines exhibited reduced susceptibility to *R. solanacearum* than the wild type plants at 7 dpi (Fig. 2B). In addition, a higher level of HR cell death displayed by darker trypan blue staining and hypersensitive response related H2O2 accumulation displayed by darker staining of e 3,3’-diaminobenzidine tetrahydrochloride (DAB) were found in leaves of *CaPti1-OE-2* and *CaPti1-OE-5* lines after *R. solanacearum* inoculation, compared with WT plants (Fig. 2C). Consistently, a lower level of *R. solanacearum* growth was found in *CaPti1-OE-2* and *CaPti1-OE-5* lines than in the wild type plants at 2 dpi (Fig. 2D). In addition, significant pronounced upregulations of *NbHSR203* (Chappell et al., 1997), *NbHSR515* (Czernic et al., 1996), *NbPR1* (Brogue et al., 1991)*, NbHIN1* (Pontier et al., 1999) were found in *CaPti1-OE-2* and *CaPti1-OE-5* lines challenged with *R. solanacearum*, compared with the wild type plants (Fig. 2E). As *CaPti1* was significantly upregulated in guard cells of pepper leaves inoculated with *R. solanacearum* by root (Supplemental Fig. S2D), we postulated that it might be involved in stomatal regulation in response to *R. solanacearum*. Therefore, the stomatal apertures in *CaPti1-OE* and WT lines with *R. solanacearum* inoculation were investigated. Upon *R. solanacearum* inoculation via root irrigation, plants of *CaPti1-OE* lines exhibited more pronounced stomatal closure compared with the WT plants (Fig. 2 F and G). Taken together, the results suggest that *CaPti1* acts positively in pepper immunity against *R. solanacearum* partially by stomatal regulation.

**Figure 2.**
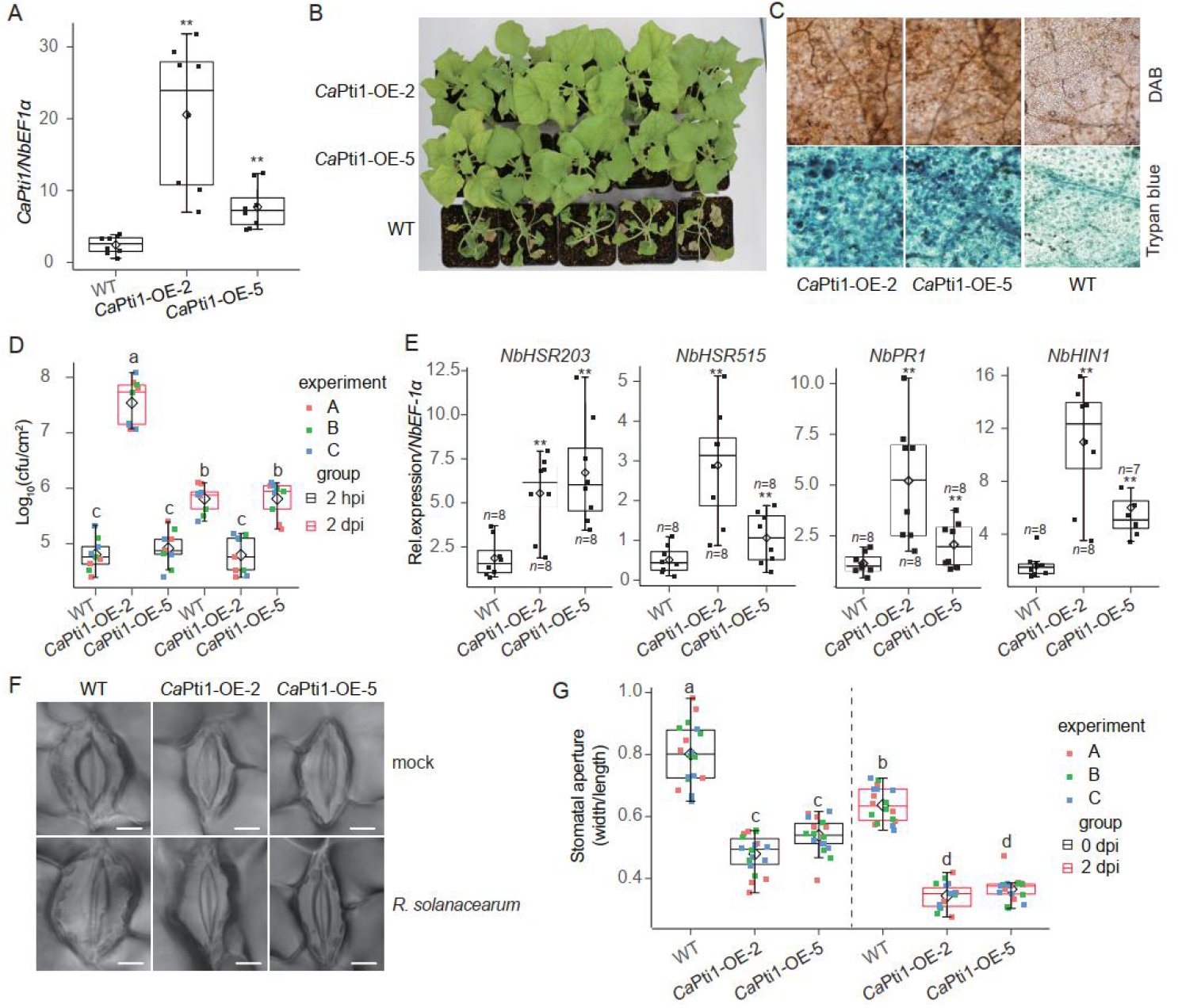
Ectopic overexpression of *CaPti1* enhanced the resistance of *N. benthamiana* plants to RSI. A, Transcript levels of *CaPti1* in *CaPti1* overexpressing lines (#2 and #5) and wild-type (WT) by quantitative RT-PCR. The transcript levels of *CaPti1* were normalized to those of *CaACTIN*. Red dots in boxplots represent 8 biological replicates (*n*=8). B-C, Bacterial symptoms (B) and DAB/trypan blue staining (C) of *R. solanacearum* inoculated eight-week-old *N. benthamiana* plants at 7 dpi. D, *R. solanacearum* growth in *R. solanacearum* inoculated *N. benthamiana* plants at 2 hours post inoculation (hpi) and 2 days post inoculation (dpi). Dots with different colors in boxplots represent 9 biological replicates (*n*=9), each comprising 6 leaf discs. Data are from 3 independent experiments. E, The transcript levels of *NbHSR203, NbHSR515, NbPR1* and *NbHIN1* at 2 dpi in *R. solanacearum* inoculated *N. benthamiana* plants. The transcript levels of each tested immunity related genes were normalized to those of *NbEF1*α. Red dots in boxplots represent at least 7 biological replicates (*n*≥7). Three technical replicates for each biological sample were used. F-G, Guard cells (F) and stomatal apertures (G) at 2 dpi in leaves of *R. solanacearum* inoculated *N. benthamiana* plants. Representative images were taken under a confocal microscope and the stomatal apertures were measured, bars = 5 μm. Dots with different colors in boxplots represent 3 independent experiments with 6 biological replicates in each experiment (*n*=18). (A, D, E and G) Asterisks indicate significant difference as determined by two-tailed *t* test (\*\**p*<0.01), and different letters indicate significant differences, as analyzed by Fisher’s protected LSD test (*p*<0.05).

### *Ca*Pti1 positively regulated dehydration tolerance through stomatal closure in an ABA signaling dependent manner

One of the mechanisms by which *R. solanacearum* destroys plants is that it infects from the roots and blocks the vascular bundles, thereby causing the aerial part of the plants to die due to lack of water (Mansfield et al., 2012). The data that *CaPti1* was upregulated by exogenously applied ABA, its induction in guard cell upon *R. solanacearum* inoculation, and its positive regulation of stomatal closure upon *R. solanacearum* inoculation indicate that *Ca*Pti1 might couple pepper immunity and drought tolerance through stomata closure probably in an ABA signaling dependent manner. To test this speculation, the effect of exogenously applied ABA, which has been believed as stomata opening regulator, on stomatal aperture was assessed in *CaPti1*-silencing and control pepper plant. Exogenously applied ABA induced a closed stomata phenotype in control pepper plants, while the stomata remained opened in TRV:*CaPti1* plants (Fig. 3, A and B), indicating that stomata closure mediated by ABA is *CaPit1* dependent. In addition, plants of *TRV:CaPit1* pepper lines exhibited larger stomatal apertures in their leaves in response to *R. solanacearum* compared with control lines after 2 dpi (Fig. 3, C and D). TRV:*CaPit1* pepper lines exhibited more sensitive to dehydration stress with lower survival rate than the control plants when they withheld watering for 15 days (Fig. 3, E and F). The plants of TRV:*CaPit1* pepper lines had lower level of fresh leaf weight, which displays water retention capacity, than control plants under dehydration condition at all of the tested time-points (Fig. 3G). Consistently, the transcripts of dehydration tolerance related genes, including *CaOSM1* and *CaOSR1*, were also analyzed after dehydration stress treatment in pepper leaves. The transcript levels of the tested genes were significantly lower in TRV:*CaPti1* plants than in the control plants (Fig. 3H). By contrast, the *CaPti1* overexpressed *N. benthamiana* exhibited enhanced tolerance to dehydration stress than the wild type control plants (Supplemental Fig. S6A), and the detached rosette leaves lost less water in *CaPti1* overexpression lines than in the WT plants (Supplemental Fig. S6B).The positive regulation of *Ca*Pti1 in the transgenic *N. benthamiana* plants was found to relate to ABA signaling, since *CaPti1* overexpressed *N. benthamiana* exhibited higher levels of seedling establishment and higher germination upon exposure to treatment of 1 μM ABA (Supplemental Fig. S6, B and C). Seeds of *CaPti1* overexpression lines and the growth of their primary roots which was more flourishing were less sensitive to ABA application than the wild type control plants (Supplemental Fig. S6, E and F). All these data suggest that the dehydration stress tolerance mediated by *Ca*Pti1 is related to stomatal closure regulated by ABA signaling.

**Figure 3.**
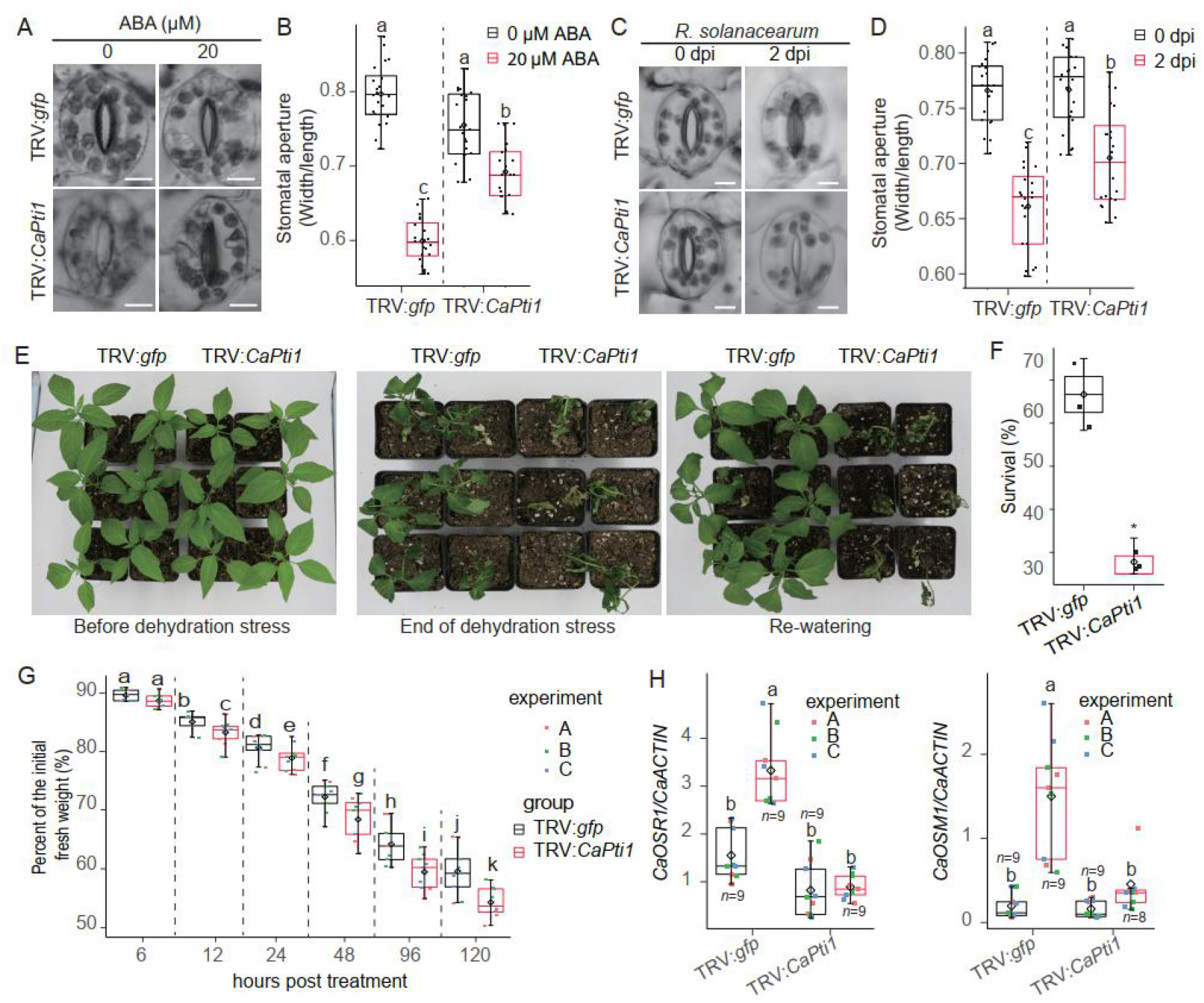
Sensitivities of pepper plants to ABA and dehydration stress were enhanced by *CaPti1* silencing. A and B, Phenotype (A) and stomatal apertures (B) of guard cells in *CaPti1*-silenced and control pepper plants after ABA treatment. Leaves treated with 0 or 20 μM ABA for 1 h by spraying were obtained from 4-week-old plants of each line. Representative images were taken under a microscope (bars= 5 μm), and the stomatal apertures were measured. The data points (*n*=21) are present in boxplots from 3 independent experiments, each calculating 7 guard cells per genotype. C-D, Guard cells (C) and stomatal apertures (D) in leaves of *CaPti1*-silenced and control pepper plants at 2 dpi after infection with *R. solanacearum* by root, bars= 5 μm. The data points (*n*=21) are present in boxplots from 3 independent experiments, each calculating 7 guard cells per genotype. E, The dehydration phenotype of *CaPti1*-silenced and control pepper withheld watering for 15 d. F, Percentage of surviving plants at 2 d post re-watering. The data points (*n*=3) are present in boxplots from 3 independent experiments, each calculating 20 plants per genotype. Asterisks indicate significantly difference, as determined by two-tailed *t* test *(*p*<0.05). G, Leaf water loss from the leaves was assessed at various time-points after leaf detachment. Dots with different colors in boxplots represent 3 independent experiments with 3 biological replicates in each experiment (*n*=9). H, Relative expression levels of dehydration responsive genes in the leaves of control and *CaPti1*-silenced pepper plants at 6 h after dehydration stress treatment. The transcript levels of indicated genes were normalized to those of *CaACTIN*. Dots with different colors in the boxplots represent at least 8 biological replicates from 3 independent experiments (n≥8). (B, D, G and H) Different letters above the bars indicate significant differences, as analyzed by Fisher’s protected LSD test (*p*<0.05).

### *Ca*ERF4 interacted with *Ca*Pti1

As a putative TyrKc domain-containing protein, *Ca*Pti1 might fulfill its function and transmits the defense signaling downward by interacting with its substrates. To isolate its possible substrates, a yeast-two-hybrid (Y2H) assay was performed using *Ca*Pti1 as a bait to screen a cDNA library of pepper plants challenged with *R. solanacearum* through root irrigation. Five positive clones were acquired (Supplemental Table S1), among which a clone encoding an ethylene responsive factor (ERF) was selected for further study. The clone shared highest sequence identity to *At*ERF4 among all of the ERFs in Arabidopsis, therefore we named it as *Ca*ERF4. The *Ca*Pti1-*Ca*ERF4 interaction was further confirmed using a one-by-one Y2H assay (Fig. 4A), *in vivo* co-immunoprecipitation (co-IP) assay via transient expression assay in *N. benthamiana* plants (Fig. 4B), pull-down (Fig. 4C), luciferase complementation imaging (LCI) assay (Supplemental Fig. S7A) as well as bimolecular fluorescence complementation (BiFC) assay (Supplemental Fig. S7B). The fluorescence signal observation in BiFC indicated that this interaction occurred in the nucleus (Supplemental Fig. S7B).

**Figure 4.**
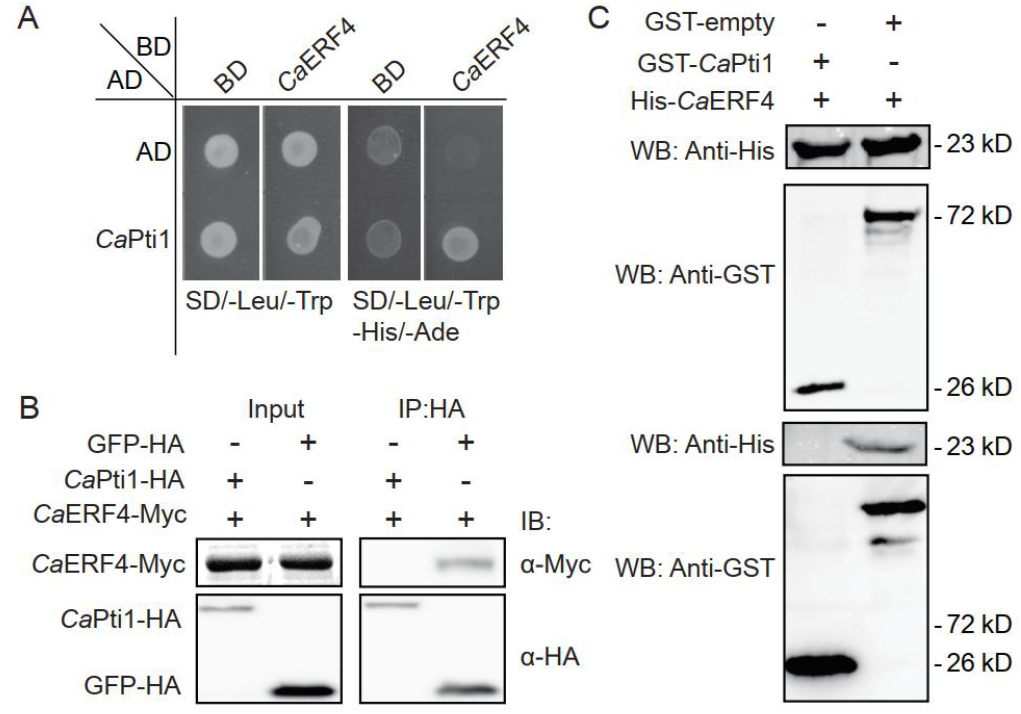
Interaction between *Ca*Pti1 and *Ca*ERF4. A, The Y2H system was used to determine the interaction of *Ca*Pti1 and *Ca*ERF4. Yeast cells were plated on SD/-Leu/-Trp/-His medium containing 50 mM 3-amino-1,2,4-triazole. B, *Ca*Pti1 and *Ca*ERF4 interaction as confirmed by co-IP. *Agrobacterium* with *Ca*Pti1-HA (GFP-HA) and *Ca*ERF4-Myc was co-infiltrated into *N. benthamiana* leaves. The leaves were harvested at 2 dpi after infiltration, and total proteins were extracted for western blotting analysis. GFP-HA was used as a negative control. C, *Ca*Pti1 physically interacts with *Ca*ERF4 *in vitro* as indicated by a pull-down assay. GST-*Ca*Pti1 or GST-bound resins were incubated with purified His-*Ca*ERF4. Co-precipitation of His-*Ca*ERF4 with the GST-binding resins was examined by western blotting using an anti-His antibody. Anti-GST antibody was used to detect GST-empty and GST-*Ca*Pti1 proteins.

To further measure the kinase activity of *Ca*Pti1 using *Ca*ERF4 as a substrate, we purified *Ca*Pti1-GST (using *Ca*CDPK2 as positive control) protein and combined it with a purified His-*Ca*ERF4 protein, which was treated with ATP to generate *in vitro* kinase reaction mixture. SDS-PAGE gels were used to separate the mixture then blotted using phosphoserine/ threonine antibody (Munoz et al., 2018). The results showed that the positive control *Ca*CDPK2 protein produced the expected band (Supplemental Fig. S7C). However, there was no indication of a label associated with any protein with a similar size to *Ca*ERF4-His or to *Ca*Pti1-GST. The results indicate that *Ca*Pti1 protein cannot phosphorylate itself or *Ca*ERF4.

### *Ca*ERF4 bound to the GCC box

It has been frequently reported that ERF proteins act as transcription factors (TF) by binding to the GCC- or DRE box elements in their target promoters (Fujimoto et al., 2000; Chakravarthy et al., 2003; Tournier et al., 2003; Liu et al., 2006; Van der Does et al., 2013). To test if *Ca*ERF4 can bind these *cis*-elements, an electrophoretic mobility shift assay (EMSA) was performed to test the DNA-binding ability of *Ca*ERF4 using Cy5-labeled double repeats of the GCC box as the probe. *Ca*ERF4 was found to form a complex with the Cy5-labeled GCC box, and the binding signal gradually decreased with the increasing addition of the unlabeled probe (cold competitor) (Supplemental Fig. S8A), indicating that *Ca*ERF4 directly bound to the GCC box *in vitro*. The trans-activation of *Ca*ERF4 in pepper leaves was examined using a β-glucuronidase (GUS) reporter assay system. A reporter vector containing GUS controlled by the *CaMV35S* core promoter (−46 to +8 bp) with two copies of GCC box in its proximal upstream region (*2xGCC-p35S_core_:* GUS) was generated. The reporter vector was co-transformed with the effector vector (*35S:CaERF4-HA*) in pepper leaves followed by immunotblot analysis and GUS activity measurement (Supplemental Fig. S8, B and C). Pepper leaves co-transformed with the reporter and the effector vector exhibited a higher level of GUS activity than the control pepper leaves, suggesting that *Ca*ERF4 might activate GUS in a GCC box-dependent manner.

### *CaERF4* acted positively in pepper immunity against *R. solanacearum*

To study the role of *Ca*ERF4, we first measured its transcriptional response to *R. solanacearum*, dehydration stress and exogenous application of ABA by RT-qPCR. *CaERF4* were found to significantly induce at transcriptional level not only by *R. solanacearum* inoculation (also in guard cells), also by exogenously applied ABA as well as by dehydration stress (Supplemental Fig. S9). By subcellular localization assay, *Ca*ERF4-GFP protein was found to accumulate exclusively in the nuclei against *R. solanacearum* (Supplemental Fig. S10), implying that *CaERF4* might be involved in pepper defense against *R. solanacearum* and tolerance to dehydration stress. To validate this speculation, *CaERF4*-silenced pepper plants and *CaERF4* overexpressed *N. benthamiana* lines were developed. The success of the gene silencing and its specificity was measured by RT-qPCR (Supplemental Fig. S11A). Similar to *CaPti1, TRV:CaERF4* pepper plants exhibited stunted growth, fewer leaves, and shorter root in pepper plants compared with the wild type control (Supplemental Fig. S11B), and exhibited more severe bacterial wilting symptoms than the control plants at 6 dpi (Fig. 5A, supplemental Fig. S11C). This susceptibility was coupled with significant increase in *R. solanacearum* growth (Fig. 5B) and high disease indices from 5 to 11 dpi *R. solanacearum* (Supplemental Fig. S11D), but coupled with lower transcript levels of defense-associated genes including *CaPR1, CaHIR1*, and *CaPO2* (Fig. 5C). Similarly, *CaERF4*-silencing significantly compromised the stomata closure (Fig. 5, D and E) and up-regulation of dehydration tolerance associated *CaOSM1* and *CaOSR1* in the distal pepper leaves (systematic response) in pepper plants inoculated with *R. solanacearum* by root (Fig. 5F). By contrast, *CaERF4*-overexpressed *N. benthamiana* plants exhibited reduced susceptibility to *R. solanacearum* via root irrigation (Fig. 5, G and H), and lower levels of bacterial multiplication (Fig. 5I), as well as more pronounced stomatal closure (Fig. 5, J and K) in response to *R. solanacearum* than the wild-type lines. All these data indicate that *Ca*ERF4 acts positively in pepper immunity against *R. solanacearum* in a way related to stomatal regulation.

**Figure 5.**
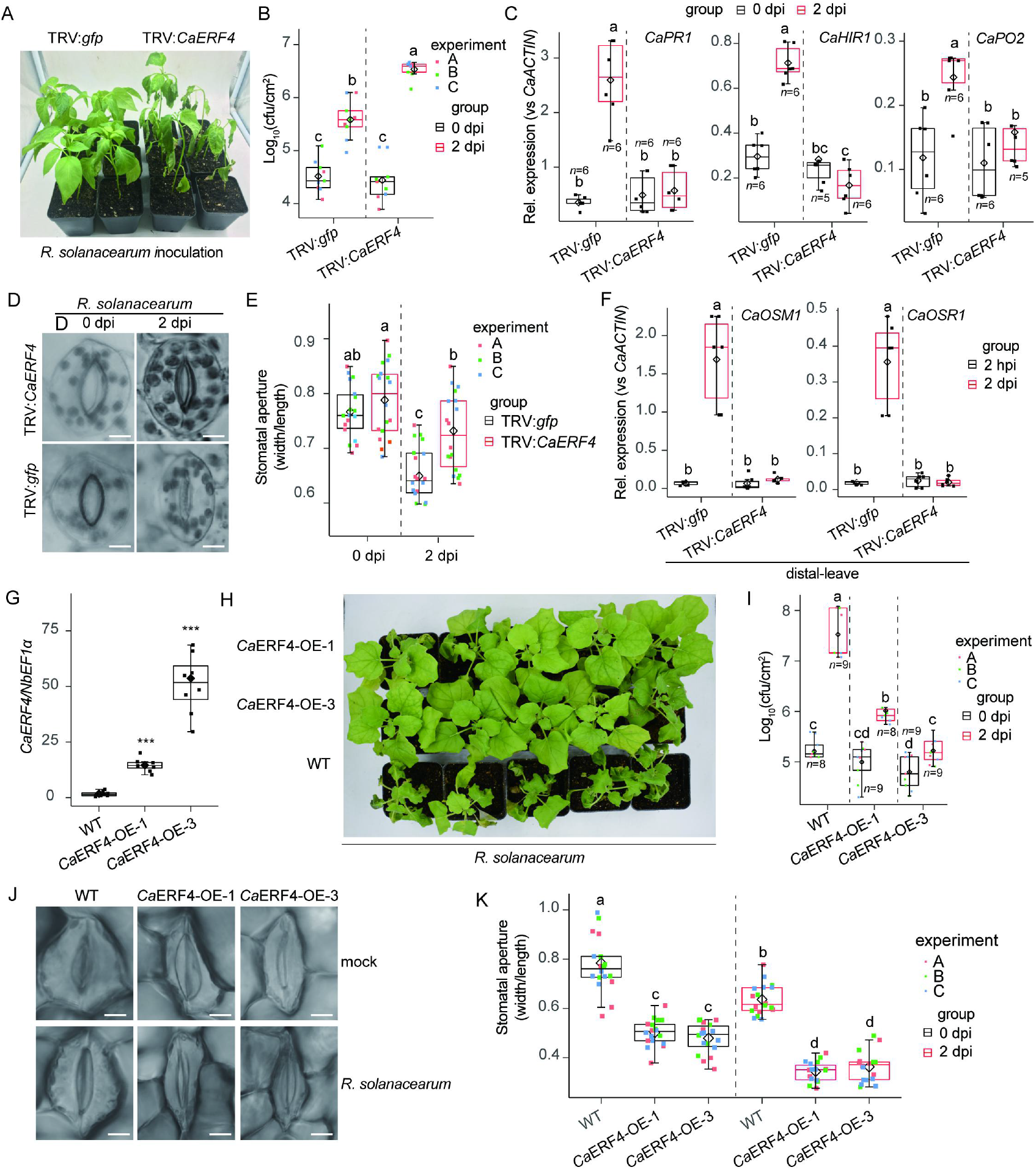
*CaERF4*-silencing compromised disease resistance of pepper plants and its overexpression enhanced disease resistance of NB plants. A, *CaERF4*-silenced pepper plants were more susceptible to the host pathogen *R. solanacearum* than the wild type plants when inoculated at roots. B, Bacterial growth in *R. solanacearum* inoculated pepper plants at 2 hpi and 2 dpi. Dots with different colors in boxplots represent *n*=9 biological replicates from 3 independent experiments. C, The expression levels of defense-related *CaPR1, CaHIR1*, and *CaPO2* in roots of *R. solanacearum* inoculated *CaERF4*-silenced and control pepper plants at 2 dpi. The transcript levels of indicated genes were normalized to those of *CaACTIN*. Dots with different colors in boxplots represent at least 5 biological replicates (*n*≥5). Three technical replicates were performed for each biological sample. D-E, Guard cells (D) and stomatal apertures (E) in leaves of *CaERF4*-silenced and control pepper plants at 2 dpi after inoculation with *R. solanacearum* at roots. *n*=18 biological replicates from 3 independent experiments. F, The transcript levels of dehydration tolerance related marker genes in distal leaves at 0 and 2 dpi after *R. solanacearum* inoculation at roots. The transcript levels of indicated genes were normalized to those of *CaACTIN*. Dots with different colors in boxplots represent *n*=6 biological replicated. Three technical replicates were performed for each biological sample. G, The transcript levels of *CaERF4* in *CaERF4* overexpressing *N. benthamiana* lines (lines 1 and 3), which were normalized to those of *NbEF1*α. Red dots in boxplots represent eight biological replicates (*n*=8). H, The phenotype of *CaERF4* overexpressing *N. benthamiana* plants challenged with RSI at 6 dpi. Asterisks indicate significant differences, as determined by the two-tailed *t* test (\**p*<0.05). I, The growth of *R. solanacearum* in *N. benthamiana* plants challenged with RSI at 2 hpi and 2 dpi. Dots with different colors in boxplots represent at least eight biological replicates from 3 independent replicates (*n*≥8). J-K, Phenotype of guard cells (J) and stomatal aperture (K) in leaves of *N. benthamiana* plants inoculated with (2 dpi) at roots or without *R. solanacearum*, bars = 5 μm. K, Dots with different colors represent 3 independent experiments with 6 biological replicated in each experiment (*n*=18). (B, C, E, F, I and K) Different letters indicate significant differences (*p*<0.05), as determined by Fisher’s protected LSD test.

### *Ca*ERF4 acted positively in response of pepper plants to dehydration stress

To test whether *Ca*ERF4 phenocopys with *Ca*Pti1 in pepper tolerance in response to dehydaration stress, we assay the effect of *CaERF4* silencing or overexpression on pepper tolerance to dehydration stress. We found that upon dehydration stress, TRV:*CaERF4* pepper plants exhibited a severe dehydration-sensitive phenotype with fully wilted leaves (Fig. 6A), lower survival rate and faster transpiration water loss compared with the control pepper lines (Fig. 6, B and C). Consistently, dehydration tolerance related *CaOSM1* and *CaOSR1* were found to be downregulated by *CaERF4* silencing (Fig. 6D). By contrast, the *CaERF4* overexpression enhanced tolerance dehydration stress (Fig. 6E), and the detached rosette leaves from overexpression lines showed significantly higher water retention capacity than did those of wild-type lines (Fig. 6F). These data indicate that *Ca*ERF4 acts positively in response of pepper plants to dehydration stress.

**Figure 6.**
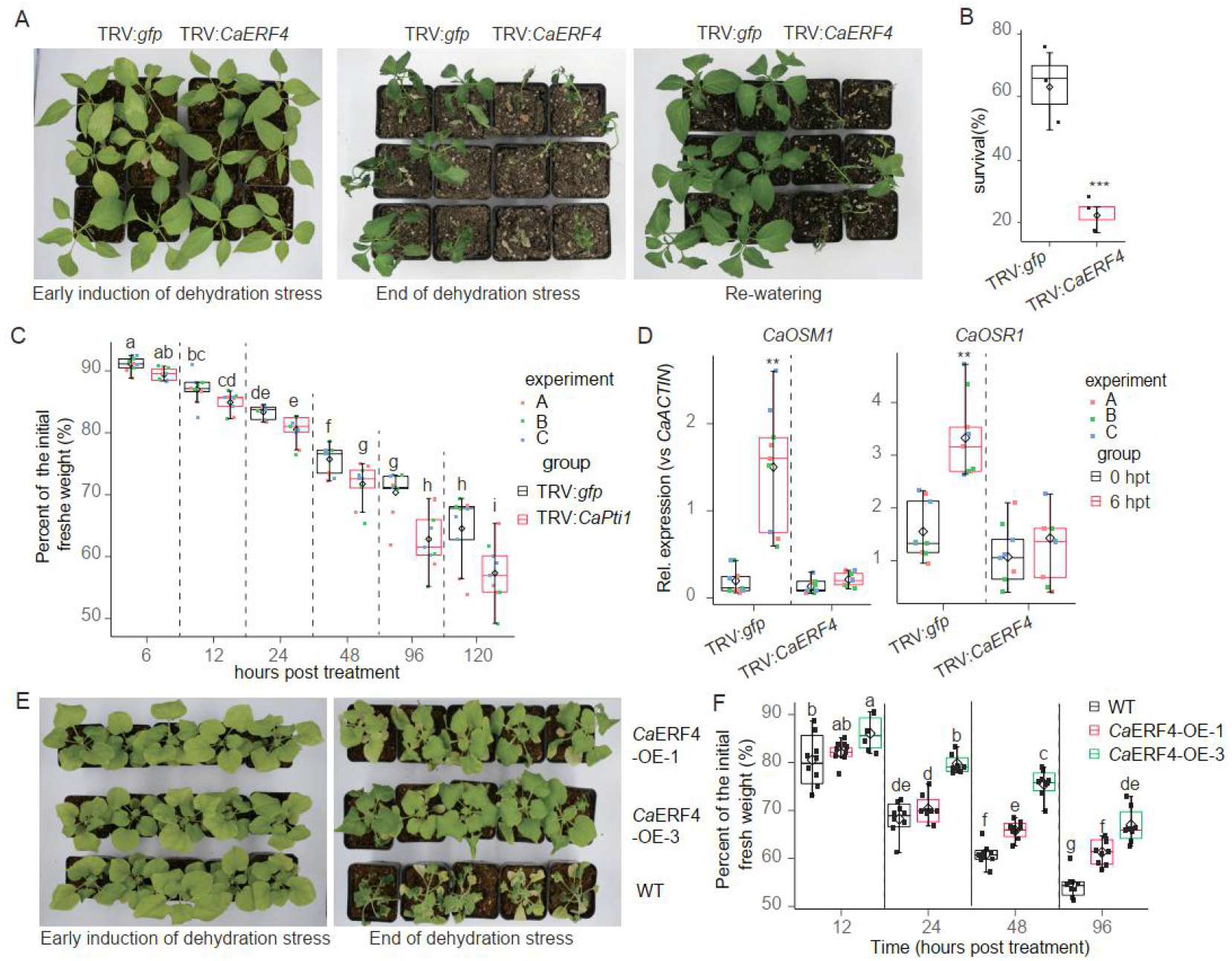
*Ca*ERF4 acted positively in dehydration stress tolerance. A, *CaERF4* silencing enhanced the sensitivity of pepper plants to dehydration stress, pepper plants were grown in pots for six weeks under well-watered conditions, and then watering was withheld for 15 days. Representative images were taken before (left) and after (middle) dehydration stress and 2 d after re-watering (right). B, Survival rates at 2 d post re-watering. Red dots in boxplots represent 3 independent experiments, each evaluating 20 plants per genotype. Asterisks indicate a significantly difference, as determined by the two-tailed *t* test (\*\*\**p*<0.001). C, Water losses from the leaves of pepper leaves at various time points after detachment. *n=9* biological replicates from 3 independent experiments. Asterisks indicate significant differences, as determined by the two-tailed *t* test (\**p*<0.05). D, The expression levels of dehydration responsive genes in the leaves of control and *CaERF4*-silenced pepper plants at 6 h post dehydration treatment. The transcript levels of indicated genes were normalized to those of *CaACTIN*. Dot with different colors in boxplots represent 3 independent experiments with 3 biological replicated in each experiment (*n*=9). Asterisks indicate significant differences, as statistically analyzed by the two-tailed *t* test (** *p* < 0.01. E, Phenotype of *CaERF4*-overexpressed *N. benthamiana* plants after withhold watering for 15 days. F, Water loss of detached *N. benthamiana* leaves at indicated time-points after detachment. *n=9* biological replicates from 3 independent experiments. In (C) and (E), the different letters indicate significant differences, as determined by Fisher’s protected LSD test (*p*<0.05).

### Transient co-expression of *CaPti1* and *CaERF4* induced more intensive cell death and stronger defense response

To assay the possible effect of *Ca*Pti1/*Ca*ERF4 interaction on defense response, we performed *Agrobacterium*-mediated transient overexpression system, which have been frequently used in pepper immunity related study (Choi et al., 2012). The success of transient overexpression of the two proteins *Ca*Pti1-HA and *Ca*ERF4-Myc were assayed by immunoblotting (Supplemental Fig. S12A). The transient overexpression of *Ca*Pti1 or *Ca*ERF4 alone triggered slight hypersensitive cell death (displayed by darker trypan blue staining), ion leakage, H2O2 accumulation (displayed by darker DAB staining), and nitric oxide (NO) accumulation in pepper leaves (Supplemental Fig. S12, B-E). However, their co-transient overexpression triggered more intensive hypersensitive cell death (Supplemental Fig. S12, B and C), higher level of ion leakage (Supplemental Fig. S12D), and higher level of H_2_O_2_ (Supplemental Fig. S12E) or nitric oxide (NO) accumulation (Supplemental Fig. S12E). Consistently, the co-transient overexpression of *Ca*Pti1 and *Ca*ERF4 also triggered the highest levels of transcript of defense-related *CaPR1* and *CaHIR1* as well as dehydration tolerance related *CaOSM1* and *CaOSR1* than that by transient overexpression of *Ca*Pti1 or *Ca*ERF4 alone (Supplemental Fig. S12F).

### *Ca*Pti1 stimulated the transcription factor activity of *Ca*ERF4

As *CaPR1, CaOSM1*, and *CaOSR1* can be regulated by *Ca*ERF4 (Fig. 5C, Fig. 6D, Supplemental Fig. S12F), we speculate that some of these genes might be directly targeted by *Ca*ERF4 and indirectly regulated by *Ca*Pti1 during pepper response to *R. solanacearum*. To test this possibility, we performed chromatin immunoprecipitation PCR (ChIP-PCR) using chromatins isolated from pepper leaves originally infiltrated with *Ca*ERF4-HA and subsequently challenged with or without *R. solanacearum*. Enrichments of *Ca*ERF4 on the promoters of *CaPR1, CaOSM1*, and *CaOSR1* were assayed by qPCR using immunoprecipitated DNA as template. Five pairs of specific primers (Supplemental Table S2) designed to amplify a group of overlapping DNA fragments covering the upstream region of the tested promoters (2,000 bp) were used to identify the enrichment of *Ca*ERF4 on the tested promoters. We found the highest enrichment of *Ca*ERF4 on the fragment C, D, D of promoter of *CaPR1, CaOSM1* and *CaOSR1* in response to *R. solanacearum*, respectively (Supplemental Fig. S13, A-C). However, these enrichments were significantly decreased in *CaPti1*-silenced pepper plants, indicating that *CaPR1* was directly targeted by *Ca*ERF4 in a *Ca*Pti1 dependent manner.

We also performed electrophoretic mobility shift assay (EMSA) to confirm the possible binding of *CaPR1, CaOSM1*, and *CaOSR1* promoters by *Ca*ERF4 *in vitro* (Fig. 7, A and B). To do this, His-*Ca*ERF4 protein was expressed in *E. coli* and purified (Supplemental Fig. S14), and several Cy5-labelled probes from the 30-bp segments were biosynthesized. The Cy5-labelled GCC1 probe from the *CaPR1* promoter (*pCaPR1*) led to a significant shift (Fig. 7C). The upper shifts efficiently competed with a 100-fold excess of unlabeled wild-type probe GCC1. However, the mutated competitor GCC1-mt exhibited no competitiveness. Nevertheless, the Cy5-labelled probe containing the GCC2 from the *pCaPR1* only caused a slight shift, compared with the GCC1 (Fig. 7C). EMSA assay also showed that *Ca*ERF4 bound to GCC1 and GCC2 in the *CaOSM1* promoter without any difference in their affinity intensities (Fig. 7D). Moreover, slight binding was detected between the Cy5-labelled DRE box in the *CaOSR1* promoter and purified *Ca*ERF4 protein (Fig. 7D). ChIP-qPCR was further conducted to investigate the effect of *Ca*Pti1 on the recruitment of *Ca*ERF4 protein to *CaPR1, CaOSM1*, and *CaOSR1* promoters; the result showed that the transient overexpression of *Ca*Pti1 increased enrichment of *Ca*ERF4 in the promoters of all of the tested immunity- or dehydration-associated marker genes (Supplemental Fig. S15A).

**Figure 7.**
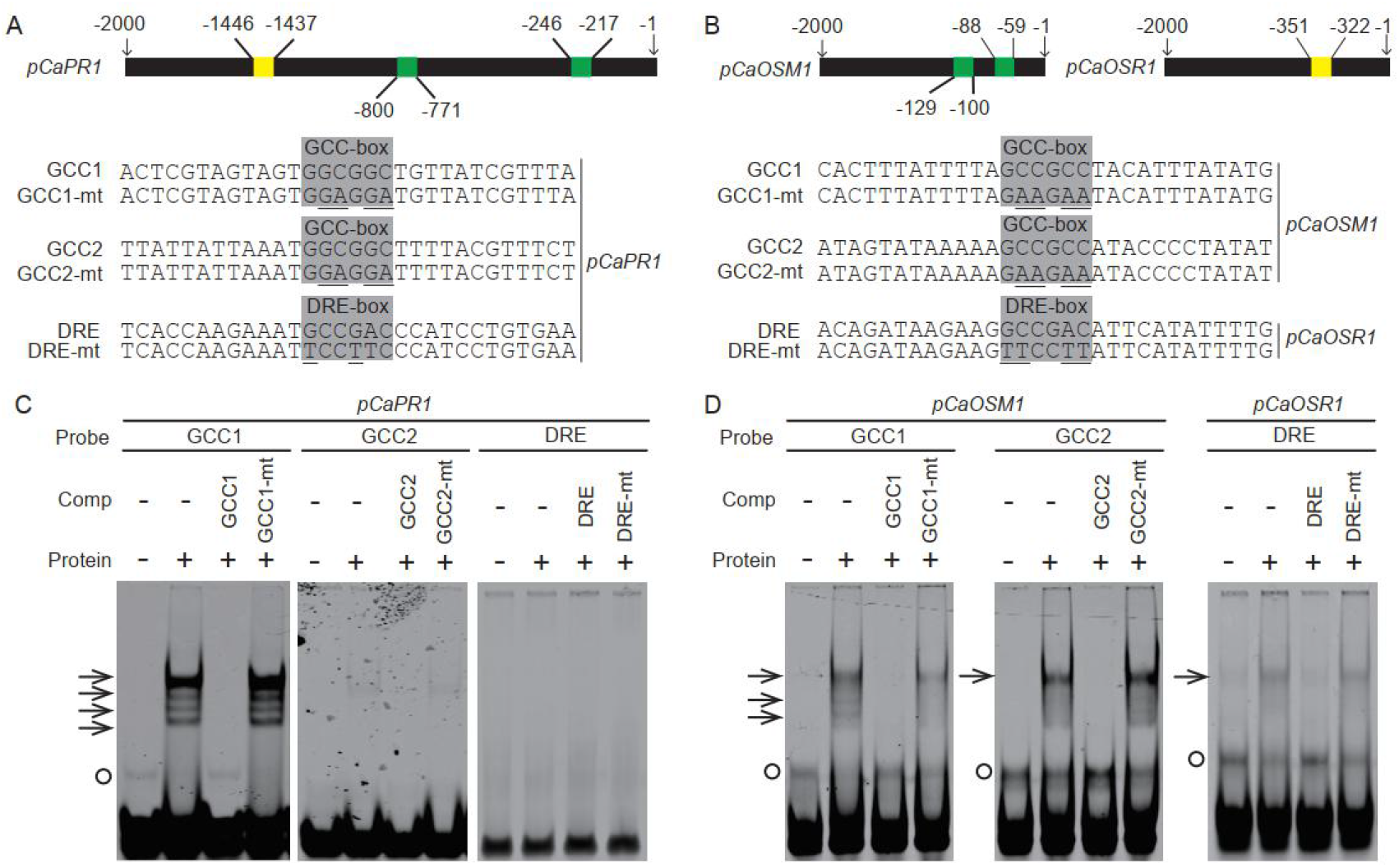
*Ca*ERF4 bound the promoters of the tested defense- or dehydration-tolerance related genes by electrophoretic mobility shift assay. A and B, The distribution of GCC or DRE box in the promoters of *CaPR1* (A), *CaOSM1* and *CaOSR1* (B). C-D, EMSA showed that the purified His-*Ca*ERF4 proteins bound the Cy5 labeled probes. Fifty-fold excess of unlabeled probes was used as competitors. Black arrows indicate specific shifts, and the circles indicate non-specific shifts or free probe.

To further test whether and how the regulation of *CaPR1, CaOMS1*, and *CaOSR1* by CaERF4 is modulated by *Ca*Pti1, we employed agroinfiltration based transient expression using effector and reporter vector with the GUS reporter gene being driven by the native promoters of the three genes. To do this, *CaPR1, CaOMS1*, and *CaOSR1* promoters were isolated from the pepper genomic DNA and fused to the β-glucuronidase (GUS) reporter gene to generate reporter vector (*pCaPR1*:GUS, *pCaOSM1:GUS*, and *pCaOSR1:GUS*). The effector vectors (*Ca*ERF4-Myc and *Ca*Pti1-HA) were also constructed. The *Agrobacterium* cells containing these vectors were co-infiltrated into pepper leaves, and the GUS activity was examined at 2 dpi. The result showed that that GUS expression driven by the tested promoters was slightly induced by *Ca*ERF4 transient overexpression alone, but the GUS activity induced by *Ca*ERF4 was enhanced, for example, 2.2-fold for *pCaPR1*, 2.6-fold for *pCaOMS1*, and 3.4-fold for *pCaOSM1*, by transient overexpression of *Ca*Pti1 (Supplemental Fig. S15B), indicating that the regulations of the tested genes by *Ca*ERF4 are potentiated by *Ca*Pti1.

All these data indicate that during pepper response to *R. solanacearum* inoculation or drought stress, *Ca*ERF4 perform its function by targeting and regulating different immunity or drought tolerance related genes, which are potentiated by *Ca*Pti1 in a protein-protein interaction manner.

## Discussion

### Dehydration tolerance and immunity against *R. solanacearum* are coupled and positively regulated by *Ca*Pti1 and *Ca*ERF4 module related to ABA signaling

Bacterial wilt is caused by *R. solanacearum* through vascular system blockade (Lowe-Power et al., 2018; MacIntyre et al., 2020), thus plant dehydration tolerance could also contribute to its resistance bacterial wilt. However, whether and how plants orchestrate immunity and defense response to dehydration stress is currently unclear. This study established that, upon *R. solanacearum* inoculation, dehydration tolerance and immunity against *R. solanacearum* are coupled and positively regulated by *Ca*Pti1 and *Ca*ERF4 in a way at least partially related to ABA signaling. Our data showed that *CaPti1* and *CaERF4* were similarly upregulated by *R. solanacearum* or exogenously applied ABA. They phenocopied each other in promoting pepper resistance to bacterial wilt not only by activating *CaHIR1, CaPO2* and SA signaling dependent *CaPR1*, but also by activating dehydration tolerance related genes, including *CaOSM1* and *CaOSR1* and by closing the stomata in an ABA dependent manner, as stomata in leaves of both *Ca*Pti1 and *Ca*ERF4 silenced pepper plants were unable to closed, by contrast, their overexpressions closed the stomata, consistent to role of ABA signaling as positive regulator in plant tolerance to dehydration stress via stomata regulation (Assmann, 2003; Chater et al., 2013; Munemasa et al., 2015; Assmann and Jegla, 2016; Yoshida and Fernie, 2018; Li et al., 2020). We speculate that the stomata closure mediated by both *Ca*Pti1 and *Ca*ERF4 is induced not for preventing *R. solanacearum* from entering, as *R. solanacearum* invade plants exclusively form the roots (Lowe-Power et al., 2018; MacIntyre et al., 2020), instead, for reducing the loss of water caused by transpiration. It is worth pointing out that the dehydration-related *CaOSR1* was activated in the leaves but not in the roots, indicating that the dehydration tolerance is mainly required for the aerial parts of the plants, which might be in water shortage due to vascular system blockade by proliferation of *R. solanacearum* (MacIntyre et al., 2020). However, how signaling mediated by ABA and that mediated by SA are coordinated upon *R. solanacearum* remain to elucidated, since antagonism have been found previously between them (de Torres-Zabala et al., 2007; de Torres Zabala et al., 2009; Moeder et al., 2010; Robert-Seilaniantz et al., 2011). It can also be speculated that both *Ca*Pti1 and *Ca*ERF4 locate downstream of ABA signaling, since their transcriptions were upregulated by exogenously applied ABA, and their overexpressions in *N. benthamiana* plants all mimicked exogenous application of ABA in closing the stomata, but the exogenously applied ABA failed to rescue the stomata closure resulted from their silencings (Figs. 3 and 5). Our data also indicate that dehydration stress response mediated by *Ca*Pti1 or *Ca*ERF4 upon *R. solanacearum* are shared by response of pepper to drought, since both of them are also upregulated at transcription level by drought stress, and their overexpression all enhanced tolerance of *N. benthamiana* plants to dehydration stress.

Importantly, *Ca*Pti1 and *Ca*ERF4 were also found to act positively in regulation of pepper growth, even though the immune or dehydration defense responses they activated are resource cost (Supplemental Figs. 4 and 11). Similar phenomenon was observed in the transcription factor IPA1 which promotes both yield and immunity in rice by sustaining a balance between growth and immunity through conditional activation and deactivation (Wang et al., 2018). To decipher the molecular mechanisms for the possible conditional activation or deactivation of *Ca*Pti1 and *Ca*ERF4 between growth and immunity, further study is required.

### *Ca*ERF4 directly and simultaneously regulate immunity and dehydration tolerance-related genes via GCC box or DRE box during pepper response to *R. solanacearum*

ERFs have been extensively studied to identify their functions in disease resistance and abiotic stress response via transcriptional regulation of their target genes in GCC box (Gutterson and Reuber, 2004; Lee et al., 2004) or DRE box, respectively (Lee et al., 2004). The loss-of and gain-of-function assays showed that *R. solanacearum* inoculation or dehydration stress induced both *CaPti1* and *CaEFR4* transcripts. Moreover, *CaEFR4* positively regulated pepper response to *R. solanacearum* by upregulating immunity-related marker genes, including *CaPR1, CaHIR1*, and *CaPO2. Ca*ERF4 also positively regulated pepper response to dehydration, indicating that *Ca*ERF4 is also involved in response to dehydration tolerance probably in a context-specific manner. Importantly, after *R. solanacaerum* inoculation, *Ca*ERF4 also upregulated *CaOSR1*, which positively regulates plant response to dehydration stress (Park et al., 2016). This suggests that transcriptional regulation of *Ca*ERF4 partially confers the activation of dehydration-related genes by *R. solanacearum*. *Ca*ERF4 can bind to GCC boxes in *CaPR1* and *CaOSM1* promoters and to DRE box in *CaOSR1* promoter, similar to *Ca*ERFLP1 (Lee et al., 2004). Notably, the affinity strengths of *Ca*ERF4 proteins binding to the two GCC boxes in the *CaPR1* promoter were significantly different, suggesting that flanking sequences can affect the *Ca*ERF4-GCC box binding. These findings indicate that *Ca*ERF4 positively regulates pepper resistance to bacterial wilt disease by directly activating immunity-related and dehydration tolerance-related genes via the binding to GCC box or DRE box.

### The targeting and regulation of *Ca*ERF4 on immunity and dehydration tolerance related genes during pepper response to *R. solanacearum* is promoted by *Ca*Pti1 by physical interaction

*Ca*ERF4, as a transcription factor, achieves its function by mediating massive transcriptional reprogramming upon pathogen perception in the nuclei, probably via upstream defense signaling integration from the cytoplasm. This study indicated that *Ca*ERF4 could be regulated at the post-translational level by interacting with *Ca*Pti1, which is upregulated by *R. solanacearum* and might locates mainly in the plasma membrane and cytoplasm on non-stressed conditions and translocate into nuclei after its being activated by upstream signaling initiated by perception of the invading pathogens such as *R. solanacearum* through R proteins such as Pto. We speculate that the nuclear *Ca*ERF4 might be activated by upstream signaling via interaction with *Ca*Pti1, thereby promoting targeting and regulating of *Ca*ERF4 to the tested immunity and dehydration tolerance related genes, similar to previous results that kinases-like MPK4 modified the targeting of WRKY33 to camalexin biosynthetic gene PAD3 (Chi et al., 2013) and ASR3 in regulation of MAMP triggered immunity (Li et al., 2015). However, we did not detect the kinase activity of *Ca*Pti1, suggesting that *Ca*Pti1/*Ca*ERF4 interaction did not lead to phosphorylation of *Ca*ERF4 without *R. solanacearum* inoculation. Nonetheless, the interaction promoted the enrichments of *Ca*ERF4 to the promoters of immunity- and drought tolerance-related genes, including *CaPR1, CaOSM1* and *CaOSR1*, thereby promoting their transcript levels. One explanation is that *Ca*Pti1 might activate *Ca*ERF4 via phosphorylation, and the kinase activity of *Ca*Pti1 is dependent on its context-specific activation by upstream signaling components, such as Pto. Besides, Pto could be activated upon recognition of effectors or PAMPs derived from pathogens (Balmuth and Rathjen, 2007; Xiao et al., 2007). In our experiment, the prokaryotic *Ca*Pti1 was not triggered by upstream signaling components; thus, no *Ca*Pti1-triggered phosphorylation of *Ca*ERF4 was detected.

Collectively, our data indicate that upon the challenge of *R. solanacearum, Ca*Pti1 is upregulated, activated and translocates from cytoplasm into the nuclei, where it promotes *Ca*ERF4 in its targeting and regulating the immunity and dehydration tolerance related genes, thereby enhancing pepper resistance to the attack of *R. solanacearum* (Fig. 8). Our results provide a new perspective for deciphering the mechanisms underlying plant resistance to bacterial wilt and its control.

**Figure 8.**
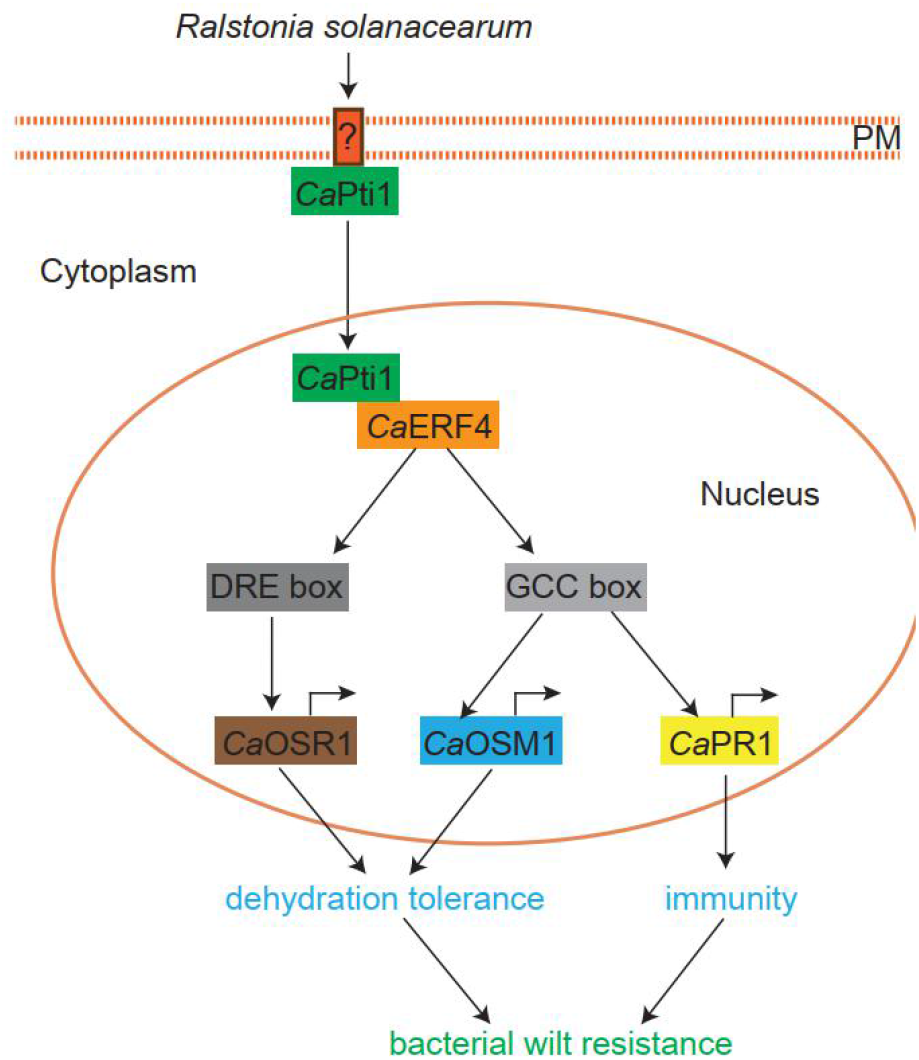
The proposed model for determining the *Ca*Pti1 and *Ca*ERF4 complex role in defense signaling and dehydration tolerance in pepper plants. Upon the challenge of *R. solanacearum, Ca*Pti1 is upregulated and activated and translocates from cytoplasm into the nuclei, where it promotes *Ca*ERF4 in its targeting and regulating the immunity and dehydration tolerance related genes, thereby enhancing pepper resistance to the attack of *R. solanacearum*.

## Materials and Methods

### Plant materials and pathogen inoculation

Seeds of pepper (*Capsicum annuum* cv. *yanshan01*) and *N. benthamiana* were germinated in sterilized ddH_2_O. The seedlings were transferred in a plastic pot and grown in a greenhouse at 25 °C, 70% relative humidity, and a 16 h photoperiod. At six weeks, the roots of fully expanded pepper and *N. benthamiana* plants were damaged by inserting a knife into the soil thrice for pathogen inoculation. The soil was irrigated with 10 ml of 10^8^ cfu/ml *R. solanacearum* FJC100301 suspended in sterilized ddH_2_O (Dang et al., 2013). The *R. solanacearum*-infected and control plants were grown in a greenhouse at 28 °C.

### Exogenous ABA and dehydration treatment

Leaves of 6-week-old pepper seedlings were separately sprayed with 100 μM ABA for phytohormone treatment. The ABA-treated pepper leaves were harvested at corresponding time-points for total RNA extraction and the reverse transcription assay as previously described (Liu et al., 2015).

Dehydration treatment assay was performed as described by Kaundal *et al*.(Kaundal et al., 2017). Briefly, VIGS and control pepper plants, control and transgenic *N. benthamiana* plants were subjected to dehydration stress treatment by not watering for 15 days. The drought tolerance was quantitatively determined by measuring the survival rate and transpiration water loss. The rosette leaves were detached from pepper plants and placed in Petri dishes to determine transpiration water loss. The plates were maintained in a growth chamber at 40% relative humidity, and fresh weight loss was determined at various time points.

### *N. benthamiana* plant transformation

The transgenic *N. benthamiana* plants were obtained following the method described (Muller et al., 1987). Briefly, GV3101 cells containing various vectors (*Ca*Pti1-OE and *Ca*ERF4-OE) were used to infect *N. benthamiana* leaf discs. The potential *N. benthamiana* transgenic plants were screened on MS medium supplemented with 500 mg L^-1^ carbenicillin and 75 mg L^-1^ hygromycin. PCR was performed to identify the positive transformants using specific primers. The plants were then self-pollinated to acquire the seeds of T1 lines and screened on 75 mg L^-1^ hygromycin-containing medium during germination. The resulting plants were self-pollinated again to obtain the seeds of T2 lines. Plants of T3 lines were selected on antibiotic medium and used for functional analysis.

### *Agrobacterium*-mediated transient expression

*Agrobacterium*-mediated transient expression was done as described with slight modification (Choi et al., 2012). Open reading frames of *CaERF4* and *CaPti1* were amplified using the primers listed in Supplemental Table S3 to generate the *Ca*ERF4-Myc and *Ca*Pti1-HA overexpression constructs. The fragments were recombined into pEarleyGate 203 (Myc tag) or pEarleyGate 201 (HA tag). *Agrobacterium tumefaciens* strain GV3101 with the constructs were cultured overnight in Luria-Bertani (LB) liquid medium containing 75 μg mL^-1^ kanamycin and 75 μg mL^-1^ rifampicin. Bacterial cells were harvested via centrifugation and were resuspended in an induction medium (10 mM MgCl_2_, 10 mM MES, pH 5.7, 200 μM acetosyringone). The resuspended bacterial cells were infiltrated into pepper and *N. benthamiana* leaves using a needleless syringe.

### VIGS assay

Virus-induced gene silencing (VIGS) assay was performed as described with slight modifications (Liu et al., 2002). Specific fragments of *CaPti1* and *CaERF4* were amplified using gene-specific primers (Supplemental Table 3) and cloned into the VIGS vector TRV2 to generate *TRV2:CaERF4* and *TRV2:CaPti1*. BLASTN in pepper genome platform (http://passport.pepper.snu.ac.kr/?t=PGENOME) was used to determine the fragment specificity. Transformed *Agrobacterium* cells with TRV1, TRV2:*CaERF4*, TRV2:*CaPti1* were grown, harvested, and resuspended in the infiltration medium (10 mM MgCl_2_, 10 mM MES, pH5.7, 200 μm acetosyringone). *Agrobacterium* cells harboring TRV1 and TRV2 were mixed at a 1:1 ratio and incubation at 25 °C for 3-4 hours, and then infiltrated into the cotyledons of pepper plants (at four-leaf-stage). The silenced and control pepper plants were used for further experiments 3-4 weeks post agroinfiltration.

### Subcellular localization

*Agrobacterium tumefaciens* strain GV3101 harboring 35S:*CaERF4-GFP, 35S:CaPti1-GFP, Ca*Pti1_pro_:*CaERF4-GFP, Ca*Pti1_pro_:*CaERF4-GFP* or 35S:*GFP*, were cultured in LB medium. The pellets were collected via centrifugation, suspended in the induction medium (10 mM MgCl_2_, 10 mM MES, pH 5.7, 200 μM acetosyringone), and adjusted to an OD595 of 0.8. The suspensions were vacuum-infiltrated into the leaves of 6-week-old *N. benthamiana* plants using a needleless syringe and maintained in the greenhouse. For 35S:*CaERF4-GFP* and *35S:CaPti1-GFP* constructs, the *N. benthamiana* leaves were subjected to fluorescent signal detection using a confocal laser scanning microscope SP8 (Leica, Wetzlar, Germany) at 48 h post infiltration. For *Ca*Pti1_pro_:*CaERF4-GFP* and *Ca*Pti1_pro_:*CaERF4-GFP* constructs, the agroinfiltrated *N. benthamiana* plants were inoculated with *R. solanacearum*, followed by fluorescence detection at 24 hpi. The emission and excitation wavelengths were 488 nm and 510-520 nm, respectively. The detached leaves were immersed in 4’, 6-diamidino-2-phenylindole (DAPI) staining solution (0.1% DAPI dissolved in 5% dimethylformamide [DMF]) for nuclei visualization. The emission and excitation wavelengths for DAPI fluorescent images were 405 nm and 430-480 nm, respectively.

### Ion conductivity measurement

Ion conductivity was measured as previously described (Liu et al., 2015). Briefly, the leaf discs were cut, washed, and incubated in 10 ml double distilled water at room temperature for one hour. A conductivity meter (Mettle toledo, Zurich, Switzerland) was then used to determine the ion conductivity.

### Histochemical staining

DAB and trypan blue staining were performed as described by Hwang (Hwang et al., 2014) with slight modifications. The detached pepper or *N. benthamiana* leaves were immersed in 1 mg/ml DAB staining buffer and kept at 25 °C overnight, then treated with 70% ethanol for DAB staining. For trypan blue staining, the detached leaves were boiled in lactophenol/ ethanol/trypan blue solution (10 ml lactic acid, 10 ml glycerol, 10 g phenol, 10 ml absolute ethanol, and 10 mg trypan blue dissolved in 10 ml distilled water) for 5 min, then destained in 2.5 g/ml chloral hydrate in distilled ddH_2_O.

### NO production and GUS activity measurement

The NO-sensitive dye 4,5-diaminofluorescein diacetate (DAF-2DA; Sigma-Aldrich, StLouis, MO) was used to monitor NO production. A needleless syringe was used to infiltrate leaves with 200 mM sodium phosphate buffer (pH 7.4) with 12.5 μM DAF-2DA, and then incubated in the dark at room temperature for one hour. A confocal laser-scanning microscope (Leica, Wetzlar, Germany) with an excitation wavelength of 470 nm was used to detect the fluorescence of diaminotriazolofluorescein (DAF-2T) and the reaction product of DAF-2DA with NO. The emission images at 525 nm were obtained at a constant acquisition time.

Protein extraction buffer [10% glycerol, 25 mM Tris-HCl, pH 7.5, 150 mM NaCl, 1 mM EDTA, 1% Triton X-100, 10 mM DTT, 1× complete protease inhibitor cocktail (Sigma-Aldrich, StLouis, MO), and 2% (w/v) polyvinylpolypyrrolidone] was used to extract total proteins from pepper leaves transiently expressing the reporter vectors to quantitatively measure GUS activity. A microplate reader (Biotek, Vermont, USA) was used to determine the rate of p-nitrophenol (γ=415 nm) release for GUS activity measurement.

### Guard cell isolation and RNA extraction

Guard cell enrichment was performed as described by (Misra et al., 2015; Kaundal et al., 2017) with slight modifications. *R. solanacearum* infected pepper leaves (10 g) without the main veins were harvested and blended thrice with 100 ml of cold, sterile water supplemented with the transcription inhibitors (0.01% cordycepin and 0.0033% actinomycin D) for 30 s. This served to inhibit the potential induction of transcript during the guard cell enrichment (Kaundal et al., 2017). The blended mixture was filtered through a 200 μm nylon mesh, and then washed using cold distilled water until the flow-through was free of mesophyll cells, debris, and plastids. The epidermal peels were digested with a mixture of 0.7% cellulysin and 0.025% macerozyme R10 (Yakult, Tokyo, Japan) with 0.1% polyvinylpyrrolidone 40 and 0.25% BSA in darkness for one hour while shaking at 150 rpm. The digest was then filtered through a 200 mm nylon mesh and washed using 750 ml of a cold basic solution (560 mM sorbitol, 5 mM MES, 0.5 mM CaCl2, 0.5 mM MgCl_2_, and 10 mM KH2PO4, pH 5.5) to remove broken epidermal cells. Trizol reagent was used to isolate RNA from guard cell enrichment.

### Yeast two-hybrid assays (Y2H)

The yeast bait vector pDEST22-*Ca*Pti1 was constructed, and the Y2H library screening was performed using the GAL4-base system following the manufacturer’s instruction (Thermo Scientific, Rockford, IL). The bait vector pDEST22-*Ca*Pti1 was co-transformed into yeast strain MaV203 with the prey vector pDEST32-*Ca*ERF4. Co-transformants were first selected on synthetic dropout (SD) agar medium without Trp and Leu (SD/-Leu/-Trp), then grown on SD/-Leu/-Trp/-His supplemented with 50 mM 3-amino-1,2,4-triazole (3-AT).

### BiFC analysis

*Ca*ERF4 and *Ca*Pti1 protein-coding sequences were amplified from pepper cDNA using attB adaptor-linked primers. They were then cloned into the BiFC vector pDEST-SCYNE and pDEST-SCYCE, respectively, to generate *Ca*ERF4-SCYNE, *Ca*Pti1-SCYCE, *Ca*ERF4-SCYCE, *Ca*Pti1-SCYNE. The *Agrobacterium tumefaciens* strain GV3101 with *Ca*ERF4-SCYNE and *Ca*Pti1-SCYCE (*Ca*ERF4-SCYCE and *Ca*Pti1-SCYNE) was co-infiltrated into the *N. benthamiana* leaves. The agroinfiltrated *N. benthamiana* plants were maintained in greenhouse for 36 h. A confocal laser scanner microscope was used for fluorescence signal detection. The excitation and emission wavelengths were 513 nm and 525-535 nm, respectively.

### Co-IP

The ORF of *CaPti1* and *CaERF4* with attB adaptors was amplified using the primers listed in Supplemental Table 3 and cloned into the gateway satellite vector pDONR207 to generate *Ca*Pti1-207 and *Ca*ERF4-207 for co-IP vectors construction. *CaPti1* and *CaERF4* fragments were transferred to their respective vectors *35S:6HA* and *35S:8Myc* after nucleotides confirmation via sequencing. *Agrobacterium* strain GV3101 with the constructs was co-infiltrated into *N. benthamiana* leaves. The protein extraction buffer [10% glycerol, 25 mM Tris/HCl (pH 7.5), 150 mM NaCl, 1 mM EDTA, 2% Triton X-100, 10 mM dithiothreitol, 1× Complete Protease Inhibitor Cocktail (Sigma-Aldrich, StLouis, MO), and 2% (w/v) polyvinylpolypyrrolidone] was used to extract total protein from the infiltrated leaves. The total protein was inoculated with anti-HA magnetic beads at 4 °C (Thermo Scientific, Rockford, IL) overnight. A magnet was used to collect the beads, and then the beads were washed thrice with tris-buffered saline and 0.05% Tween 20 (TBST). SDS-PAGE was used to separate the eluted proteins with different sizes, and immunoblotting was used for analysis. Anti-HA-peroxidase and anti-Myc-peroxidase antibodies were used to detect HA- and Myc-fusion proteins.

### Pull-down assay

Recombinant glutathione S transferase (GST)-*Ca*Pti1 and His-*Ca*ERF4 proteins were expressed in *Escherichia coli* (*E. coli*) cells. GST-*Ca*Pti1 and GST (empty vector) proteins were purified using glutathione resins from the *E. coli* cells and incubated with purified His-*Ca*ERF4 protein. The complex was purified using BeaverBeads™ glutathione (GSH) and separated by SDS-PAGE. Anti-GST and anti-His antibodies were used to detect GST-*Ca*Pti1 and His-*Ca*ERF4, respectively.

### Kinase assay

GST-tagged *Ca*Pti1 protein was expressed in *E. coli* cells and purified. Purified *Ca*Pti1-GST protein was then combined with a purified *Ca*ERF4 protein tagged with a His epitope tag. The combination was treated with ATP to generate *in vitro* kinase reaction mixture. SDS-PAGE gels were used to separate the mixture then blotted using phosphoserine/threonine antibody (Munoz et al., 2018).

### Electrophoretic mobility shift assay

Cy5-labeled double-stranded DNA fragments of 30 bps in length containing the GCC or DRE box in the promoter of *CaPR1, CaOSM1, CaOSR1* and their mutated version were synthesized to generate probes. Purified His-*Ca*ERF4 protein was incubated with wild type or mutated probe labeled in 5× binding buffer [200 mM Tris-HCl (pH 7.5), 375 mM KCl, 6.25 mM MgCl_2_, 25% glycerol, 1 mM DTT). The mixture was incubated on ice for 45 min and then run on a PAGE gel, followed by scanned on an Odyssey@ CLX instrument (LI-COR, Nebraska, USA).

### Quantitative real-time RT-PCR assay

qRT-PCR was performed as previously described (Liu et al., 2015). Briefly, TRIzol reagent (Thermo Scientific, Rockford, IL) was used to extract the total RNA from pepper and *N. benthamiana* plants. DNase I was used to digest the genomic DNA in the isolated RNA samples. A cDNA first-strand synthesis kit (Vazyme, Nanjing, China) was used for the reverse-transcription of total RNA (500 ng) following the manufacturer’s instructions. Bio-rad real-time PCR system (Bio-rad, CA, USA) and the SYBR premix *Ex-Taq* II system were used for qPCR analysis. The pepper *CaACTIN* or *N. benthamiana NbEF1*α were used as internal controls. The primers used in this study are listed in Supplemental Table 3.

### Immunoblot analysis and chromatin immunoprecipitation (ChIP) assay

Total proteins were extracted from the leaves as previously described (Choi et al., 2008). Equal amounts of protein were separated via SDS-PAGE and blotted to polyvinylidene fluoride membranes for immunoblot analysis. The membranes were probed with HA-, Myc- or GFP antibodies at 1:5000 dilution. Horseradish peroxidase-conjugated goat anti-rabbit IgG was used as a secondary antibody.

ChIP assays were performed as described with slight modifications (Singh and Szabo, 2012). *CaPti1*-silenced and control pepper plant leaves transiently transformed with *35S:CaERF4-HA* were harvested at 24 h post *R. solanacearum* inoculation and cross-linked in 1% formaldehyde solution for 10 min. Chromatin was isolated as described by Kaufmann *et al*. (Kaufmann et al., 2010). The resuspended chromatin was sonicated into fragments (200 to 500 bps long) at 4 °C using an ultrasonic crusher (Covaris, Massachusetts, USA). The sheared chromatin was immunoprecipitated with anti-HA antibodies (Sigma-Aldrich, StLouis, MO), washed thrice using Tris-Buffered Saline buffer supplemented with 0.5% Tween 20 (TBST) buffer, and reverse cross-linked. Finally, the acquired DNA was eluted with sterilized double-distilled H2O. The 10% non-immunoprecipitated DNA was reverse cross-linked and used as input control. Quantitative PCR was used to assess both the immunoprecipitated DNA and input DNA using specific primers listed in Supplemental table S2/3.

### Statistical analyses

The differences between two groups were indicated by single (statistically significant *p*<0.05), double asterisks (very significantly, *p*<0.01) or triple asterisks (extreme significantly, *p*<0.001), respectively (Two-tailed *t* test). The differences among multiple groups were indicated by different letters (*p*<0.01), as calculated with Fisher’s protected least-significant-difference (LSD) test.

## Accession Numbers

Sequence data from this article can be found in the EMBL/GenBank data libraries under the following accession numbers: pepper *CaPti1* (XM_016710753), *CaERF4* (XM_016725123), *CaPR1* (AAC06244.2), *CaPR2* (M60460), *CaACS1* (X65982), *CaPti1-like* (XM_016710755), *CaDEF1* (AF442388), *CaPR1* (AF348141.1), *CaPO2* (DQ489711), *CaHIR1* (AY529867), *CaABR1* (CA524559), *CaOSM1* (AY262059), *CaOSR1* (KT693385), *CaACT1* (*AY572427*); *N. benthamiana NbHSR203* (CAA54393.1), *NbHSR515* (X95342), *NbPR1b* (X66942), *NbPR3* (X51425), *NbPR4* (AF154635.1), *NbPR1a/c* (X05959), *Nb* EF1α (D63396), Arabidopsis *AtPti1* (NP_567882); *Nicotiana attenuata NaPti1* (XP_019260946.1); *Nicotiana tabacum NtPti1* (XP_016492642.1); *Solanum lycopersicum SlPti1* (XP_004252961.1); *Solanum tuberosum StPti1* (XP_006349879.1,); *Ziziphus jujuba ZjPti1* (XP_015887013.1).

## Supplemental Material

**Supplemental Figure S1.** Comparison of amino acid sequences of *Ca*Pti1 with representative related orthologs of other species.

**Supplemental Figure S2.** Expression profiles of *CaPti1* in pepper plants.

**Supplemental Figure S3.** The subcellular localization of *Ca*Pti1 in *N. benthamiana* leaves.

**Supplemental Figure S4.** The silencing efficiency detection and developmental phenotype in *CaPti1*-silenced pepper plants.

**Supplemental Figure S5.** The expression analysis of immunity and dehydration related maker genes in root and leaves of *CaPti1*-silenced and control pepper plants when inoculated with *R. solanacearum*.

**Supplemental Figure S6.** The overexpression of CaPti1 reduced the sensitivity of *N. benthamiana* plants to exogenously applied ABA in to dehydration stress.

**Supplemental Figure S7.** Interaction between *Ca*Pti1 and *Ca*ERF4 in planta and test of phosphorylation of *Ca*ERF4 by *Ca*Pti1 using anti-Phosphoserine/threonine antibody.

**Supplemental Figure S8.** DNA-Binding and transcription activation of *Ca*ERF4.

**Supplemental Figure S9.** Expression profiles of *CaERF4* in pepper plants.

**Supplemental Figure S10.** Subcellular localization of *Ca*ERF4 driven by native promoter against *R. solanacearum* inoculation.

**Supplemental Figure S11.** The developmental phenotype and silencing efficiency detection in *CaERF4*-silenced pepper plants.

**Supplemental Figure S12.** The effect of co-transient overexpression of *CaPti1* and *CaERF4* in pepper leaves on HR cell death and expression of immunity related genes

**Supplemental Figure S13.** Recruitment of *Ca*ERF4 to the promoters of immunity- and dehydration tolerance associated marker genes by ChIP-qPCR.

**Supplemental Figure S14.** Prokaryotic expression and purification of His-*Ca*ERF4 protein in *E. coli* cells.

**Supplemental Figure S15.** The regulations of the immunity- and dehydration-related genes by *Ca*ERF4 are potentiated by *Ca*Pti1.

## Acknowledgments

The authors thank Mark D. Curtis for kindly providing the Gateway destination vectors and Dr. S. P. Dinesh-Kumar of Yale University for the pTRV1 and pTRV2 vectors.

## Notes

Funding: This work was supported by the Natural Science Foundation of Fujian Province, China (2020J01532), the Excellent Youth Foundation of Fujian Agriculture and Forestry University (xjq201913), the National Natural Science Foundation of China (31501767, 30971718, and 31372061), and Development Fund Project of Fujian Agriculture and Forestry University (CXZX2018114, CXZX2020008A, and CXZX2019024G). The funders had no role in study design, data collection, data analysis, decision to publish, or preparation of the manuscript.

